# Identification of two new trichome-specific promoters of *Nicotiana tabacum*

**DOI:** 10.1101/620021

**Authors:** Mathieu Pottier, Raphaëlle Laterre, Astrid Van Wessem, Aldana M. Ramirez, Xavier Herman, Marc Boutry, Charles Hachez

**Affiliations:** Louvain Institute of Biomolecular Science and Technology, University of Louvain, 1348 Louvain-la-Neuve, Belgium

**Keywords:** Rubisco small subunit, Major Allergen Mal D 1.0501, Cembratrien-ol Synthase, Copal-8-ol diphosphate Synthase, Lipid Transfer Protein, Cytochrome P450 oxygenase

## Abstract

*Main conclusion pRbcS-T1* and *pMALD1*, two new trichome-specific promoters of *Nicotiana tabacum*, were identified and their strength and specificity were compared to those of previously described promoters in this species.

*Nicotiana tabacum* has emerged as a suitable host for metabolic engineering of terpenoids and derivatives in tall glandular trichomes, which actively synthesize and secrete specialized metabolites. However, implementation of an entire biosynthetic pathway in glandular trichomes requires the identification of trichome-specific promoters to appropriately drive the expression of the transgenes needed to set up the desired pathway. In this context, RT-qPCR analysis was carried out on wild-type *N. tabacum* plants to compare the expression pattern and gene expression level of *NtRbcS-T1* and *NtMALD1*, two newly identified genes expressed in glandular trichomes, with those of *NtCYP71D16, NtCBTS2α, NtCPS2,* and *NtLTP1,* which were reported in the literature to be specifically expressed in glandular trichomes. We show that *NtRbcS-T1* and *NtMALD1* are specifically expressed in glandular trichomes like *NtCYP71D16, NtCBTS2α,* and *NtCPS2*, while *NtLTP1* is also expressed in other leaf tissues as well as in the stem. Transcriptional fusions of each of the six promoters to the *GUS-VENUS* reporter gene were introduced in *N. tabacum* by *Agrobacterium*-mediated transformation. Almost all transgenic lines displayed GUS activity in tall glandular trichomes, indicating that the appropriate cis regulatory elements were included in the selected promoter regions. However, unlike for the other promoters, no trichome-specific line was obtained for *pNtLTP1:GUS-VENUS*, thus in agreement with the RT-qPCR data. These data thus provide two new transcription promoters that could be used in metabolic engineering of glandular trichomes.

## Introduction

Trichomes are epidermal outgrowths covering most of aerial tissues in a large number of plant species. Several types of trichomes (unicellular or multicellular, glandular or non-glandular) can be observed in a single plant species. Among those, glandular trichomes are characterized by cells forming a glandular structure that secretes or stores specialized (also called secondary) metabolites (e.g., phenylpropanoids, flavonoids, acyl sugars, methylketones, and terpenoids). Many of these possess antimicrobial and antifungal properties or act as a defense barrier against herbivorous insects (Schilmiller et al. 2008).

The specialized metabolites secreted by glandular trichomes, which might represent up to 17 % of the leaf dry weight in *Nicotiana tabacum* (tobacco), have been largely exploited over centuries (Wagner et al. 2004). One of their most ancient uses originates from their aromatic properties and fragrances. Besides, these specialized metabolites constitute an interesting source of pharmaceuticals and food additives. Some specialized metabolites are only found in a single plant species or even a single plant cultivar and often at low concentration (e.g., taxol found in *Taxus sp.*, artemisinin in *Artemisia annua* or cannabinoids in *Cannabis sativa*). Therefore, natural resources are often insufficient to reach the global need (Van Agtmael et al. 1999; Yoon et al. 2013), while the complex stereochemistry of these compounds often prevents their full chemical synthesis in a cost-effective way.

In order to increase the overall yield, metabolic engineering strategies are undertaken in a variety of host species (Kirby and Keasling 2009; Marienhagen and Bott 2013). Advances in plant biotechnology and increasing knowledge in specialized metabolism also make possible to exploit plants as production platforms. One of the main advantages of using them is that they are photoautotrophic organisms, therefore requiring simple and cheap growth conditions, which accounts for a cost-effective production (Kempinski et al. 2015). In addition, their ability to deal with membrane proteins such as P450 enzymes and posttranslational modifications such as glycosylation, are two key features frequently limiting in prokaryotic hosts (van Herpen et al. 2010).

Terpenoids and derivatives are the most abundant plant specialized metabolites in terms of sheer number and chemical diversity (for review, see Croteau et al. 2000; Bouvier et al. 2005; Gershenzon and Dudareva 2007) and *N. tabacum* has emerged as one of the most suitable plant hosts for their biosynthesis (Moses and Pollier 2013; Lange et al. 2013; Wang et al. 2016). Indeed, *N. tabacum* synthesizes a very high amount of a limited range of specialized metabolites (Huchelmann et al. 2017). This combined to its high biomass, its fast growth rate, and its easy genetic transformation make it an interesting host to implement the biosynthesis pathways of terpenoid compounds and derivatives thereof.

However, engineering terpenoid biosynthetic pathways using ubiquist promoters frequently leads to severe phenotypes including dwarfism, chlorosis, and decreased seed production due to the cytotoxicity of these compounds or detrimental impact on the biosynthesis of essential metabolites (Saxena et al. 2014; Gwak et al. 2017; reviewed in Huchelmann et al. 2017). To avoid these adverse effects, a fine control of the spatiotemporal expression of the transgenes, restricting the biosynthesis of potentially cytotoxic metabolites to specialized organs, is desirable (Huchelmann et al. 2017). Since *N. tabacum* glandular trichomes contain an important pool of terpenoid precursors and have naturally evolved to deal with high concentrations of terpenoids, they make ideal targets to develop such a metabolic engineering approach. For this purpose, identification of trichome-specific transcription promoters is required.

A proteomic comparison was recently performed in *N. tabacum* between proteins extracted from tall glandular trichomes and those extracted from other plant organs (Laterre et al. 2017). This led to the identification of 47 proteins that were more abundant in tall glandular trichomes, the most enriched ones being a putative PR-10 type pathogenesis-related protein, namely Major Allergen Mal D 1.0501 (MALD1) and a Ribulose-1,5-Bisphosphate Carboxylase/oxygenase Small subunit (RbcS-T1) (Laterre et al. 2017). For both, semi- quantitative RT-PCR supports trichome-specific localization of their corresponding transcripts (Harada et al. 2010; Laterre et al. 2017).

This suggests that *the NtMALD1* and *NtRbcS-T* promoters may confer trichome-specificity, as those of CYtochrome P450 oxygenase 71D16 (*NtCYP71D16*), Copal-8-ol diPhosphate Synthase 2 (*NtCPS2*), Lipid Transfer Protein 1 *(NtLTP1*), and CemBraTrien-ol Synthase 2α (*NsCBTS2*α) previously described (Wang et al. 2002; Ennajdaoui et al. 2010; Choi et al. 2012; Sallaud et al. 2012). However, these six genes were investigated separately, preventing one to compare their transcript levels. In addition, for some of them, cell-type specificity monitored by the GUS reporter gene was not described in other organs than leaves. The present study thus aimed at comparing the expression patterns and expression levels of *NtCYP71D16, NtCBTS2*α*, NtCPS2, NtLTP1, NtRbcS-T1*, and *NtMALD1* in *N. tabacum*. Their transcript levels in trichomes and different organs were compared. Transcriptional fusions of each promoter to *GUS-VENUS* were expressed in transgenic *N. tabacum* plants. GUS staining corroborate transcripts data and indicate that all promoters, except for *pNtLTP1*, can be used to drive trichome-specific expression.

## Materials and methods

### Plant material and plant growth conditions

*Nicotiana tabacum* cv Petit Havana SR1 (Maliga et al. 1973) plants were used in this work. For the *in vitro* cultures, seeds were sterilized by immersion in 1 ml 70% (v/v) ethanol for 1 min and then in 1 ml 50% (v/v) commercial bleach for 2 min. Seeds were then washed three times with 1 ml of sterile MilliQ water and kept at 4°C, in the dark, during 48 h for stratification. Sterilized seeds were sown on solid Murashige and Skoog (MS) medium [4.33g.l^-1^ MS salts (MP Biochemicals, Solon, OH, USA; www.mpbio.com), 3% (w/v) sucrose, 1% (w/v) agar, pH 5.8 (KOH)] and placed in the growth chamber at 25°C under a 16 h photoperiod (50 μmol photon m^−2^ sec^−1^). For the soil cultures, seeds were stratified before being sown in potting soil (DCM, Grobbendonk, Belgium; dcm-info.com). Isolated plantlets coming from potting soil or *in vitro* conditions were transferred to Jiffy pots (Gronud, Norway; www.jiffypot.com) before being transferred to bigger pots containing potting soil (DCM). Plants on soil were grown under controlled conditions, in a phytotron set at 25°C and with a 16 h photoperiod (300 μmol photon m^−2^ sec^−1^).

### Tissue isolation, RNA extraction and cDNA synthesis

Trichomes were removed from tissues of 6-week-old plants following the cold-brushing method (Wang et al. 2001). For gene expression in trichomes during leaf development, the analysis was performed in triplicate. Trichomes were isolated from leaves at different developmental stages defined here by leaf length: < 2.5 cm (stage I), between 2.5 cm and 6.5 cm (stage II), between 6.5 cm and 15 cm (stage III), and > 15 cm (stage IV). For gene expression in different tissues, the analysis was performed in three to five replicates on roots, trichomes-free stems, trichomes-free leaves, and leaf trichomes (pool of leaves from stage I to stage III) from 6-week-old plants, and flowers from 10-week-old plants. For each biological replicate (except for isolated trichomes), 100 mg of material was pre-ground in liquid nitrogen using a mortar and pestle. Pre-ground tissues and isolated trichomes were ground in 2 mL Precellys tubes containing 200 μl of ceramic beads Zirmil (0.5 mm, Saint Gobain Zipro, Le Pontet, France) and 500 μl of lysis/2-Mercaptoethanol solution of the Spectrum^TM^ Plant Total μ RNA Kit (Sigma-Aldrich, St. Louis, Missouri, USA; http://www.sigmaaldrich.com). Samples were subjected to four consecutive 30 s grinding periods at 6,000 rpm using a Precellys 24 (Bertin Technologies, Montigny-le-Bretonneux, France). The homogenates were centrifuged at 1,000 *g* for 3 min (Eppendorf 5430, Hamburg, Germany). The subsequent steps of the RNA extraction were performed on the supernatants according to the manufacturer’s specifications, except that the 56 °C incubation step was omitted. RNA was eluted in 50 µl elution buffer and quantified using a spectrophotometer (Nanodrop® ND-1000, Isogen Life Science, The Netherlands; www.isogen-lifescience.com). Genomic DNA contamination was eliminated by using the On-Column DNase I Digestion Set (Sigma-Aldrich, St. Louis, Missouri, USA; www.sigmaaldrich.com). The RNA was finally flash frozen in liquid nitrogen and stored at - 80°C. DNA-free RNA (500 µg) was used for reverse transcription using the Moloney Murine Leukemia Virus Reverse transcriptase (Promega, Madison, Wisconsin, USA; be.promega.com) and oligo(dT)_18_. Reverse transcription mixture was added according to the manufacturer’s specifications. After adding the transcriptase, samples were incubated for 5 min at 25°C, followed by 1 h at 42°C and 5 min at 85°C, placed on ice for 5 min, aliquoted, and stored at -20°C.

### Gene expression

Gene-specific RT-qPCR primers listed in Supplemental Table S1 were designed at the 3’end of the coding sequence, (size, about 100 bp; melting temperature, 60°C) using OligoPerfect™ Designer (www.thermofisher.com). cDNA (5 µl, 17 fold diluted) was used as a template in 20 µl RT-qPCR reaction, which also contained 10 µl of the Power SYBR green PCR master mix of qPCR master mix plus for SYBR Green I (Eurogentec, Seraing, Belgium, https://secure.eurogentec.com/eu-home.html) and 5 µl of primer mix (1.3 µM each). Amplification was performed on an ABI 7500 Real-Time PCR system (Waltham, Massachusetts, USA; http://www.thermofisher.com). Primer specificity was confirmed by analysis of the melting curves. For each tissue, primer amplification efficiency (≥ 95%) was determined using five standards from serial dilutions of a cDNA pool of the biological replicates used for gene expression analysis. Relative transcript levels were calculated following the 2^−ΔΔCt^ method (Livak and Schmittgen 2001) with the geometric mean of mitochondrial ATP-synthase β-subunit (*NtATP2*), ubiquitin (*NtUBQ*), and elongation factor α (*NtEF1*α) transcripts used as reference for comparison between different tissues (three to five replicates), and of *NtATP2*, *NtUBQ*, and actin (*NtACTIN*), for comparison between different leaf developmental stages (three replicates). For absolute quantification (three replicates), PCR products amplified by gene-specific RT-qPCR primers listed in Supplemental Table S1 were cloned in pGEM-T Easy vector (Promega, Madison, Wisconsin, USA) prior to their sequencing. Constructs were linearized by *PstI* restriction, purified using Nucleospin Extract II kit (Macherey-Nagel, Düren, Germany) and rigorously quantified through UV (260 nm) absorption using a spectrophotometer (Nanodrop® ND-1000, Isogen Life Science, The Netherlands; www.isogen-lifescience.com). For each quantified purified linear plasmid, the copy number was determined according to the following equation (Godornes et al. 2007): copy number = (vector amount [g]) × 6.023 × 10^23^ [molecules/mole]) / (660 [g/mole/base] × size of the vector+insert [bases]. Absolute transcript levels were then determined through the absolute standard curve method. Thus, for each studied gene, standards (2 x 10^6^, 2 x 10^5^, 2 x 10^4^, 2 x 10^3^ copies) obtained by serial dilution of the purified linear plasmids were included in duplicate in q-PCR plates used to study gene expression during trichome development.

### Generation of plants expressing *PROMOTER:GUS-VENUS* fusions

The transcription promoter regions of *NtRbcS-T1* (1993 pb; GenBank accession: MG493459.1) and *NtMALD1* (1974 pb; GenBank accession: MG493458.1) were identified blasting the EST corresponding to *NtRbcS-T1* (GenBank accession: DV157962) and *NtMALD1* (GenBank accession: FS387666) coding sequences to the genome of *N. tabacum* TN90 in the Solgenomics database (http://solgenomics.net). The promoter regions of *NsCBTS2*α (985 bp; GenBank accession: HM241151.1), *NtLTP1* (849 bp; GenBank accession: AB625593.1), *NtCYP71D16* (1852 pb; GenBank accession: AF166332.1), and *NtCPS2* (1448 bp; GenBank accession: HE588139.1) were defined as previously (Wang et al. 2002; Ennajdaoui et al. 2010; Choi et al. 2012; Sallaud et al. 2012). Promoter regions were amplified by PCR using as a template genomic DNA prepared from *N. tabacum* or *N. sylvestris* leaves and the primers listed in Supplemental Table S2. The amplified fragments were inserted in the pGEM®-T Easy Vector (Promega, Madison, Wisconsin, USA; www.promega.com) and sequenced. Cloned fragments were cleaved using *HindIII* (or *NotI* for *pNtMALD1* and *pNtRbcS-T1*) and *KpnI*, prior to their insertion in a pAUX3131 construct (Navarre et al. 2011), upstream of the *GUS-VENUS* coding sequence. The fusion construct was excised using *I-SceI* and inserted into the pPZP-RCS2-nptII plant expression vector (Goderis et al. 2002), also cut with *I-SceI*. The construct was introduced into *Agrobacterium tumefaciens* LBA4404 virGN54D (van der Fits et al. 2000) for subsequent *N. tabacum* leaf disc transformation (Horsch et al. 1986). For each construct, 24 to 45 independent transgenic lines were generated and finally transferred to soil to be analyzed by GUS staining.

### GUS histochemical analysis

Histochemical staining of plant tissues for GUS activity was conducted during 3h30 as described previously (Bienert et al. 2012). However, to allow substrate access to the trichomes covered by the oily exudate, incubation was performed in the presence of 1% Triton X-100.

### Statistical analysis

All tests were performed using the R software. For q-PCR, data were analyzed using *kruskal.test* (Kruskal–Wallis) function for multiple comparisons. For multiple comparisons, *nparcomp* package was used to perform Tukey post-hoc test when significant differences were detected (P < 0.05). Different letters indicate significant differences between samples.

## Results

In a 2D gel analysis of glandular trichome proteins from *N. tabacum*, several spots were identified as trichome-specific proteins, among which RbcS-T1 and MALD1 (Laterre et al. 2017). Here, the RNA levels of *NtRbcS-T1* and *NtMALD1* as well as of *NtLTP1*, *NtCYP71D16, NtCBTS2*α, and *NtCPS2* previously reported as genes specifically expressed in tall glandular trichomes, were compared in trichomes and different *N. tabacum* organs. To do so, leaves were frozen in liquid nitrogen and carefully scratched with a brush to collect the trichomes. RT-qPCR assays were then performed on RNA extracted from trichomes, roots, trichome-free leaves, and trichome-free stems of six-week-old plants as well as from flowers of 10-week-old plants. Unlike for leaves and stems, trichomes could not be retrieved from flower sepals and petals. Because different organs had to be compared, it was important to use reference genes, whose expression little varies according to the organ. The transcript levels of ubiquitin (*NtUBQ*), mitochondrial ATP-synthase β-subunit (*NtATP2*), actin (*NtACTIN*), and elongation factor α (*NtEF1*α), frequently described as reference genes, were thus monitored. On the basis of this analysis, we found that *NtUBQ*, *NtATP2,* and *NtEF1*α genes were the most stable, therefore the geometric mean of their transcripts was used to normalize the data (Supplemental Fig. 1). For each studied gene, the relative expression level in trichomes was arbitrarily set to one. All six investigated genes showed a higher relative expression level in isolated trichomes compared to the levels observed in roots, leaves, stems or flowers (Fig. 1). *NtCYP71D16, NtCBTS2*α*, NtCPS2*, *NtRbcS-T1*, and *NtMALD1* exhibited very low or even undetectable expression in roots, trichome-free leaves and trichome-free stems, while higher transcript levels were found for *NtLTP1* in leaves and stems. Expression was observed in flowers for the six genes but, as noted above, sepal and petal trichomes could not be removed from these organs.

**Fig. 1.**
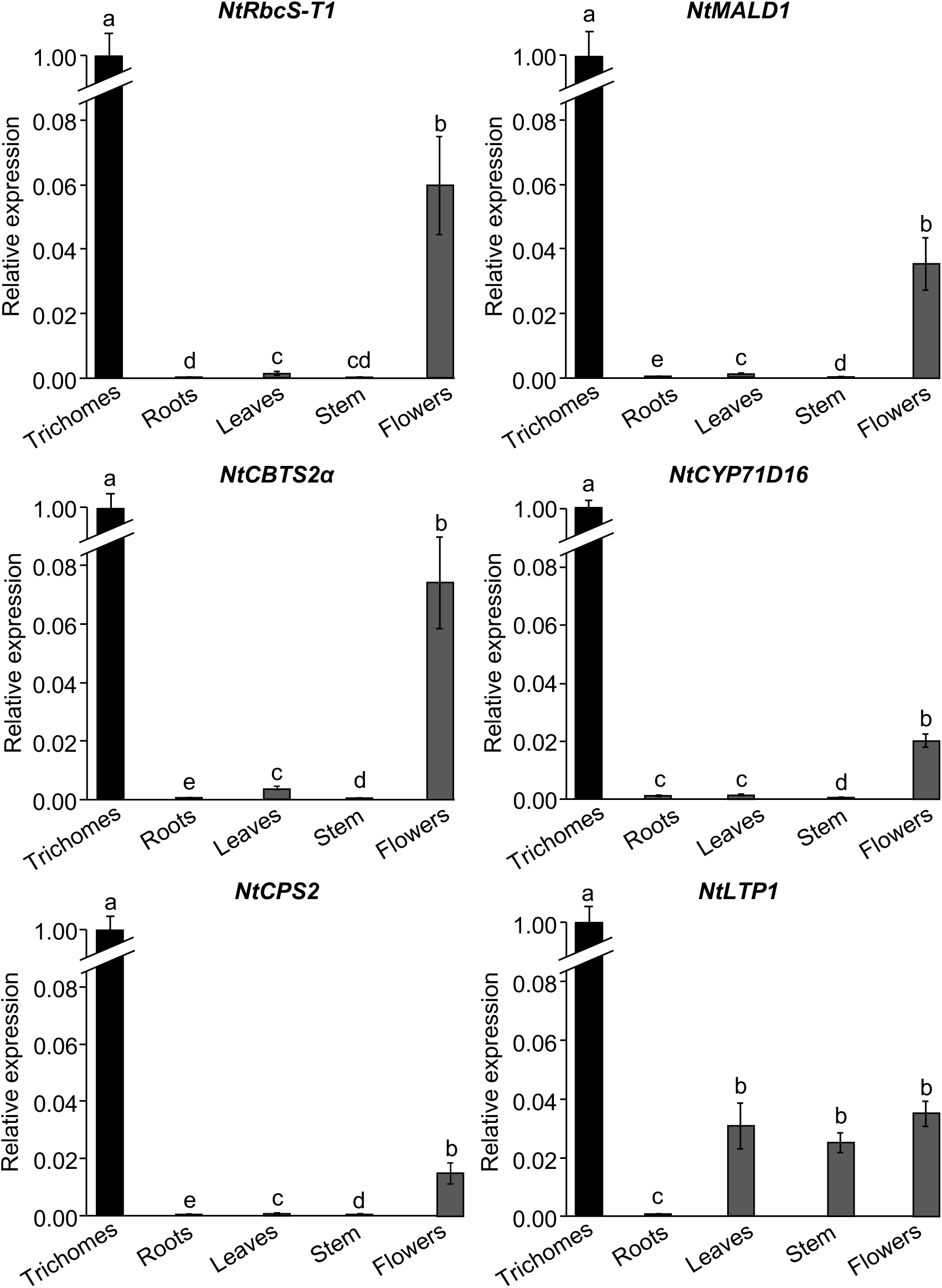
Transcript levels in different organs of *N. tabacum*. Transcript levels were normalized to the mean of those of *NtATP2* and *NtUBQ* genes. Results are shown as mean ± SD of three to five repeats. Different letters indicate significant differences according to a Kruskal-Wallis test (p < 0.05) followed by a Tukey post hoc test

As most of these genes are involved in the biosynthesis (*NtCYP71D16, NtCBTS2*α, and *NtCPS2*) or transport (*NtLTP1*) of specialized metabolites secreted by mature glands, we wondered whether the leaf developmental stage could impact their expression in trichomes. Thus, glandular trichomes were isolated from leaves at different developmental stages, arbitrarily defined by the leaf length: < 2.5 cm (stage I), between 2.5 cm and 6.5 cm (stage II), between 6.5 cm and 15 cm (stage III), and > 15 cm (stage IV). In this experiment, *NtUBQ*, *NtATP2*, and *NtACTIN* transcripts were the most stable, therefore the geometric mean of their transcripts was used to normalize the data (Supplemental Fig. 2). While the transcript level of *NtLTP1* appeared stable during leaf development, expression of the other five genes steadily increased until stage III where it reached a plateau (Fig. 2). The opposite trend was observed for elongation factor α (*NtEF1*α), which peaked at stage I (Supplemental Fig. 2), confirming that the observed increasing level of these five genes is not an artifact of the normalization method. Among them, *NtRbcS-T1* was the gene for which the transcript level increased the most with leaf development (4-fold increase). Expression of *NtCBTS2*α and *NtCYP71D16* involved in the biosynthesis of cembrenes, the major subgroup of diterpenes produced by *N. tabacum* glandular trichomes, also exhibited a large increase (3.8- and 3.6-fold, respectively) (Fig. 2). A more moderate increase was found for *NtMALD1* (2.6-fold) and *NtCPS2* (2.4-fold) transcripts, the latter being involved in the biosynthesis of another subgroup of diterpenes, namely labdanes.

**Fig. 2.**
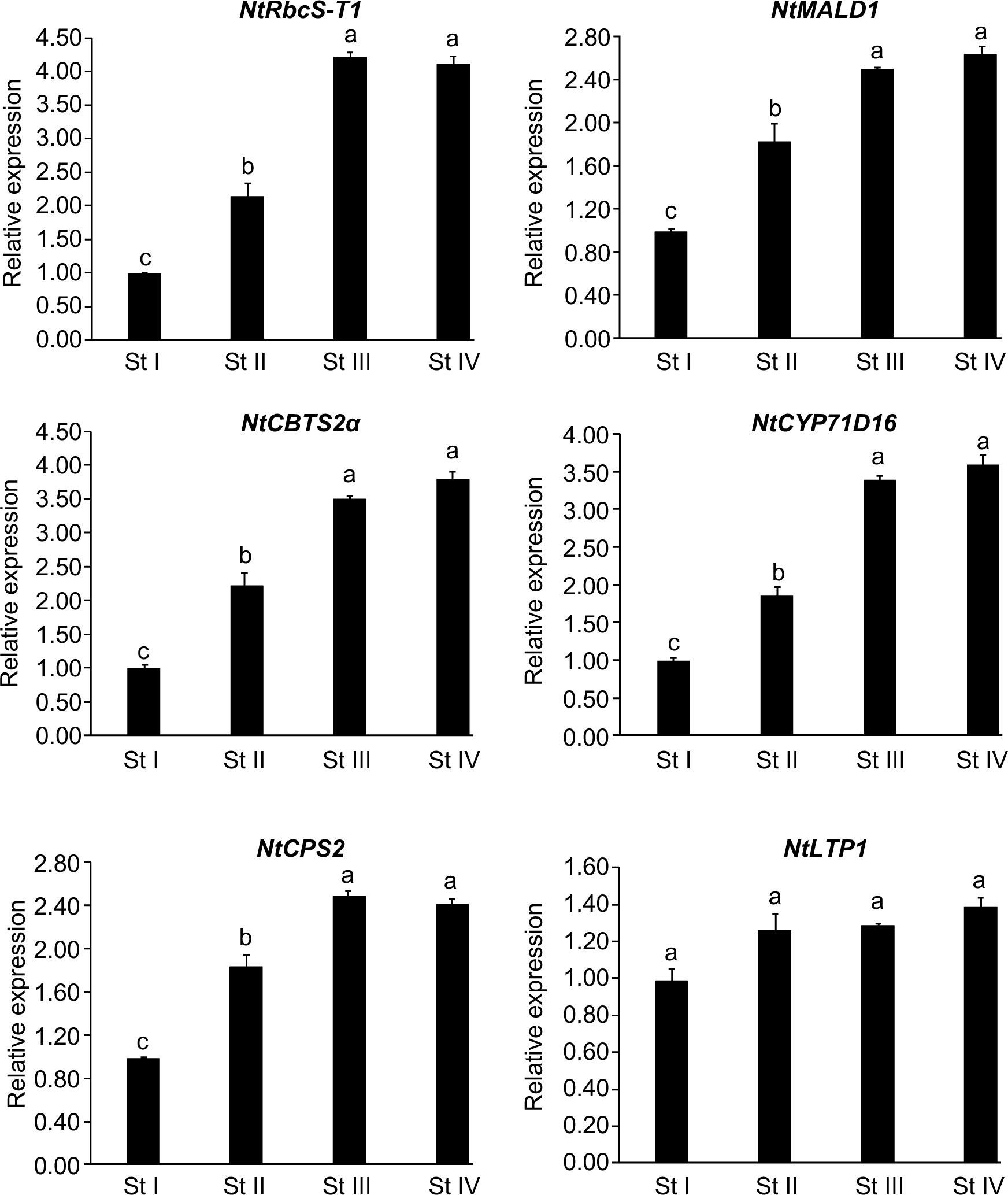
Transcript levels in trichomes isolated from *N. tabacum* leaves at different developmental stages. Transcript levels were normalized to the geometric mean of those of *NtUBQ*, *NtATP2*, and *NtACTIN* genes. St: leaf developmental stage. Stage 1: leaf length < 2.5 cm; stage II: leaf length between 2.5 cm and 6.5 cm; stage III: leaf length between 6.5 cm and 15 cm; stage IV: leaf length > 15 cm. Results are shown as mean ± SD of three repeats. Different letters indicate significant differences according to a Kruskal-Wallis test (p < 0.05) followed by a Tukey post hoc test

The absolute expression levels of all six genes of interest was then determined using the absolute standard curve method in isolated trichomes for developmental stage III at which all genes reached a maximal expression, (Fig. 3, see Material and methods for details). Several control genes, some of which were used to normalize the relative expression data shown in Figures 1 and 2, were also added to the study for comparison purposes. Among control genes, the absolute expression levels (Fig. 3) were in agreement with previously published data in other Solanaceae species (Lu et al. 2012; Lacerda et al. 2015). Genes involved in cembrene production, *NtCBTS2*α (78.0 copies/pg), and *NtCYP71D16* (67.9 copies/pg), were the most expressed genes at stage III (Fig. 3), while a lower expression was found at this stage for *NtMALD1* (40.8 copies/pg)*, NtLTP1* (28.2 copies/pg)*, NtCPS2* (labdane diterpenes, 11.1 copies/pg) and *NtRbcS-T1* (5.1 copies/pg).

**Fig. 3.**
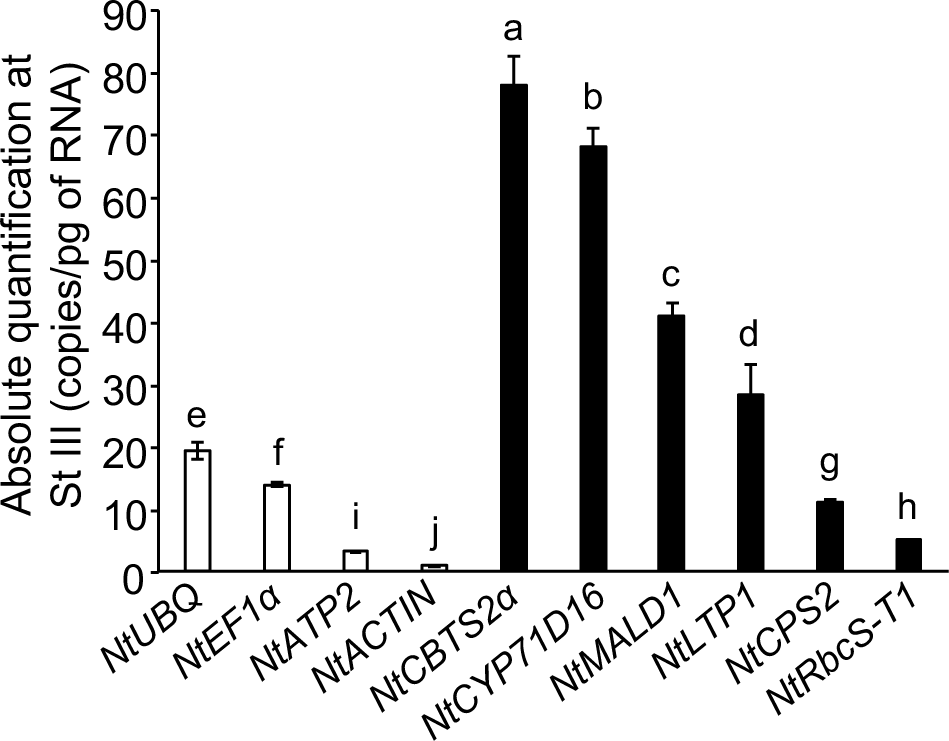
Absolute transcript levels at stage III of leaf development in *N. tabacum*. Absolute transcript levels were determined as indicated in the Material and methods. Results are shown as mean ± SD of three repeats. Different letters indicate significant differences according to a Kruskal-Wallis test (p < 0.05) followed by a Tukey post hoc test

To further confirm the trichome-specific expression pattern observed by RT-qPCR, we generated transcriptional reporter lines using a *GUS-VENUS* coding sequence. In the 2D gel analysis which led to the identification of trichome-specific proteins, two spots had been identified as trichome-specific RbcS (Laterre et al. 2017). At that time, only the *N. benthamiana* genome sequence was available and a RbcS transcription promoter (*pNbRbcS-T*) corresponding to the minor RbcS spot had been retrieved from this species and characterized (Laterre et al. 2017). Once the sequence of a *N. tabacum* genome became available, we identified *pNtRbcS-T1* (accession: MG493459.1) as the promoter of the gene corresponding to the major trichome RbcS spot (NtRbcS-T1; accession: DV157962). The *NtMALD1* promoter (accession: MG493458.1), corresponding to the NtMALD1 spot (accession: FS387666) was identified as well. The *GUS-VENUS* coding sequence was fused to *N. tabacum* genomic fragments of 1993 bp and 1974 bp upstream of the translation initiation codon of *NtRbcS-T1* and *NtMALD1*, respectively (Supplemental Fig. 3). For the other genes, the previously published promoter regions, i.e. 985 bp (*NsCBTS2*α), 849 bp (*NtLTP1*), 1852 bp *(NtCYP71D16*), and 1448 bp (*NtCPS2*) (Wang et al. 2002; Ennajdaoui et al. 2010; Choi et al. 2012; Sallaud et al. 2012) were isolated and similarly fused to the *GUS-VENUS* coding sequence. These constructs were introduced in *N. tabacum* through *Agrobacterium tumefaciens*-mediated transformation. For each construct, 24 to 45 independent transgenic lines were generated and their GUS activity was monitored in T_0_ and then confirmed on T_1_ lines. A large majority (83-95%) of the generated lines displayed GUS activity in tall glandular trichomes. This indicates that appropriate cis-sequences required for expression in tall glandular trichomes are present in the promoter sequences fused to the reporter gene. With the exception of the *pNtLTP1:GUS-VENUS* reporter (see below), between 42 and 54% of the lines for the other constructs had their GUS activity restricted to glandular trichomes on aerial parts of the plant (examples are displayed for each construct in Fig. 4a-r and Supplemental Fig. 4) while no signal was recorded in roots (Supplemental Fig. 4). In these lines, trichome expression was further confirmed by following VENUS fluorescence on living tissue by confocal microscopy (Fig. 4). The *NtCPS2* promoter provided strong expression in both leaf short and tall glandular trichomes, while the other four constructs only labelled tall glandular trichomes (Fig. 4j-l).

**Fig. 4.**
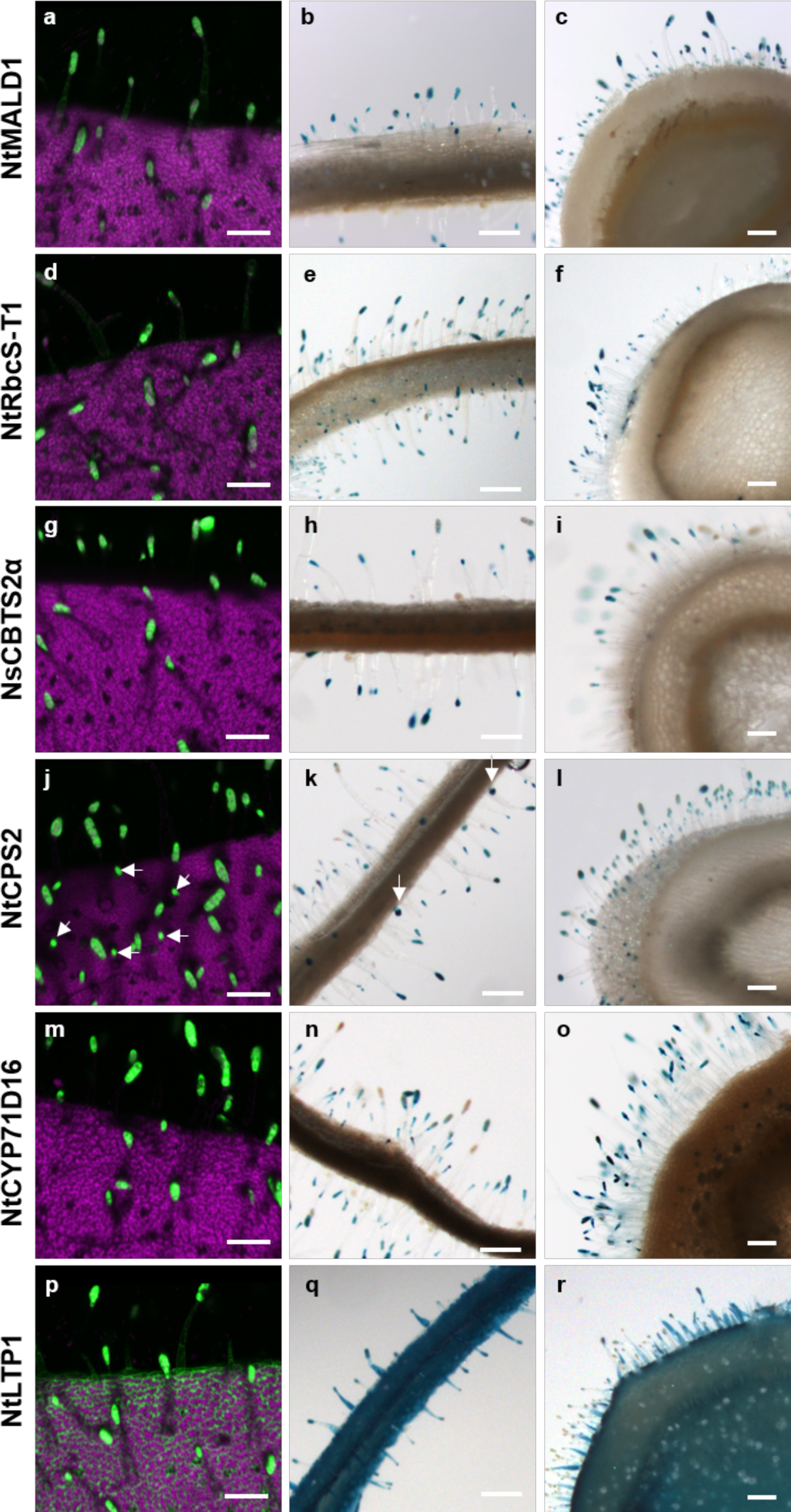
GUS-VENUS activity in leaves and stems of *N. tabacum* transgenic lines. Venus detection by confocal imaging and GUS staining were performed on transgenic lines expressing *GUS-VENUS* under the control of the promoter region of *NtMALD1* (**a**-**c**), *NtRbcS-T1* (**d**-**f**), *NsCBTS2*α (**g**-**i**), *CPS2* (**j**-**l**, white arrowheads point to the labelling of short glandular trichomes), *NtCYP71D16* (**m**-**o**), and *NtLTP1* (**p**-**r**). Left panels: 3D reconstruction of leaf tissue expressing the VENUS reporter as detected by confocal microscopy. Green: VENUS signal, magenta: chlorophyll autofluorescence. Middle panels: GUS staining in leaf cross-sections. Right panels: GUS staining in stem cross-sections. Scale bars: 200 μm

For each of these five constructs, between 46 and 58% of the lines had their expression extended to other cell types, with a pattern and an intensity varying according to the line, a typical consequence of the position effect (see discussion). As an exception, all *pNtLTP1:GUS-VENUS* lines displayed GUS activity in various leaf and stem tissues (Fig. 4p-r and Supplemental Fig. 4), confirming the absence of trichome specificity revealed by RT-qPCR data.

## Discussion

In this work, the tissue-specific expression pattern of six *N. tabacum* genes, namely *NtLTP1*, *NtCYP71D16*, *NtCBTS2*α, *NtCPS2*, *NtRbcS-T1*, and *NtMALD1*, was analyzed. In fact, although these genes were previously described as trichome-specific, their trichome-specific expression at the transcript level had not yet been quantified and compared. We performed this comparison through RT-qPCR using RNA isolated from trichomes and different plant organs as well as *GUS-VENUS* reporter genes using, when available, previously published promoter sequences.

RT-qPCR analysis showed that all these genes except for *NtLTP1*, are specifically expressed in trichomes in *N. tabacum* (Fig. 1). Regarding *NtLTP1* transcripts, they were also identified in leaf and stem tissues cleared from trichomes (Fig. 1). This observation is in line with previously published semi-quantitative RT-PCR which showed that *NtLTP1* is expressed in different organs (Fig. 1; Harada et al., 2010). Apart from *NtLTP1*, whose expression was almost constant during leaf development, that of the other five genes was lower at an early stage of leaf development and reached a maximum at stage III (Fig. 2), presumably when the specialized metabolism in which they are involved is fully operating. This is also true for *NtRbcS-T1* and this observation is in agreement with the hypothesis that in glandular trichomes, Rubisco recycles the CO_2_ released by the specialized metabolism (Pottier et al. 2018). These expression data may help choose appropriate trichome-specific promoters to drive the expression of a transgene for metabolic engineering purposes. Although *NtCYP71D16* and *NtCBTS2*α lead to higher expression level in trichomes at stage III of leaf development, *NtCPS2* and *NtMALD1* promoters should lead to a more homogenous expression of transgenes among leaves at different developmental stages.

Our analysis of *GUS-VENUS* reporter lines revealed that, in almost all of them, the six promoters drove gene expression in the head cells of tall glandular trichomes of *N. tabacum* (Fig. 4). However, in agreement with the transcript level analysis (Fig. 1; Harada et al., 2010), we observed GUS reporter activity in other organs than trichomes in all the lines expressing the *NtLTP1:GUS-VENUS* construct (Fig. 4p-r and Supplemental Fig. 4p-r). The examination of similar lines by Choi et al., (2012) has also revealed some GUS reporter activity in other cell types than trichomes while the expression in the stem was not displayed. We thus conclude that the *NtLTP1* promoter does not confer trichome-specific expression. On the contrary, trichome-specific GUS activity was observed in lines expressing any of the five other constructs (Fig. 4 and Supplemental Fig. 4), which is consistent with our RT-qPCR analysis (Fig. 1). However, other lines displayed GUS expression in trichomes, but also in other cell types, with a profile varying from line to line for a given reporter construct (data not shown). This likely results from a position effect due to the random insertion of the T-DNA in the plant cell genome. The genomic environment surrounding the integrated cassette (structure of chromatin, presence of enhancers/silencers near the insertion site) is known to alter the expression level and profile of transgenes (Kohli et al. 2010; Hernandez-Garcia and Finer 2014). Between independent lines, and thus different insertion sites, this position effect differs according to the proximal endogenous regulatory elements.

In conclusion, a key and unique feature of glandular trichomes is their ability to synthesize and secrete large amounts of a limited panel of specialized metabolites. Taking advantage of the pool of natural precursors to produce specific metabolites in glandular trichomes by metabolic engineering would therefore be of high biotechnological interest. This requires the availability of transcriptional promoters specifically active in these structures that could be used to efficiently drive the expression of the transgenes coding for the enzymes needed to implement the pathway in a cell-type specific way. In this respect, the identification of the *NtMALD1* and *NtRbcS-T1* promoters and their comparison with previously identified trichome-specific promoters are promising tools for expressing entire biosynthesis pathways in glandular trichomes of *N. tabacum*.

## Author contributions

MP, RL, CH and MB designed the experiments and analyzed the data. MP, RL, AVW, AR, XH and MB performed experiments. CH, MP and MB wrote the manuscript.

## Acknowledgments

The authors are grateful to Joseph Nader for his technical contribution. This work was supported by the Belgian Fund for Scientific Research (Grant ID: MIS – F.4522.17), the Interuniversity Poles of Attraction Program (Belgian State, Scientific, Technical and Cultural Services), and an EU Marie Skłodowska-Curie fellowship (Project ID: 658932) to MP.

## Supplementary data

**Supplemental Fig. 1** Transcript levels of control genes in different organs of *N. tabacum*

**Supplemental Fig. 2** Transcript levels of control genes in trichomes isolated from *N. tabacum* leaves at different developmental stage

**Supplemental Fig. 3** Molecular constructs used to generate transgenic *N. tabacum* expressing the *GUS-VENUS* reporter gene under the control of trichome-specific promoters

**Supplemental Fig. 4** GUS activity in various tissues of *N. tabacum* transgenic lines.

**Supplemental Table S1** List of primers used for RT-qPCR.

**Supplemental Table S2** List of primers used to amplify the promoter sequences.

