## Supplemental Tables for "Identification of two new trichome-specific promoters of *Nicotiana tabacum*"

**Supplemental Table S1** List of primers used for RT-qPCR.

| | |
|---|---|
| ***qNtCBTS2α*** | F : 5´- TTTCAAGATTGGCCGATACA -3´<br>R : 5´- TTCGACAAACAACGAAGCAA -3´ |
| ***qNtCYP71D16*** | F : 5´- AGGAGAATTTGCCCTGGAAT -3´<br>R : 5´- CCAGTTGGGAGTTTCCAGTC -3´ |
| ***qNtLTP1*** | F : 5´- CTCCTTGCGTCCCTTATTTG -3´<br>R : 5´- CAATTGCATGCAGTCTTTCG -3´ |
| ***qNtCPS2*** | F : 5´- TCCAAAATGACAAGGTGACG -3´<br>R : 5´- TTGGTTGATGTTGCTTGAGG -3´ |
| ***qNtRbcS-T1*** | F : 5´- CAGCAGCATAGAAGAAGTCAC -3´<br>R : 5´- ACTTCAACCATAAACCTTGAGG -3´ |
| ***qNtMALD1*** | F : 5´- CATGCCTGATGGAGGATGTA -3´<br>R : 5´- GCCTCAACAGCCTTGACAAT -3´ |
| ***qNtEF1α*** | F : 5´- GGACATGCGTCAAACTGTTG -3´<br>R : 5´- TTCTTCTGAGCAGCCTTGGT -3´ |
| ***qNtUBQ***[a] | F : 5´- GAGGAATGCAGATCTTCGTG -3´<br>R : 5´- TCCTTGTCCTGGATCTTAGC -3´ |
| ***qNtATP2*** | F : 5´- GGTTCCTTAGCCAGCCTTTC -3´<br>R : 5´- CCAACACTCCCTGGAAACTG -3´ |
| ***qNtACT*** | F : 5´- TTTCCTGGAATTGCTGATAGGATGA -3´<br>R : 5´- AGCCAAAATAGAACCTCCAATCCAA -3´ |

[a]Previously published (Pandey *et al.* 2014)

**Supplemental Table S2** List of primers used to amplify the promoter sequences. Restriction sites are in bold and annealing parts are in capital letters.

| | |
|---|---|
| ***pNtCYP71D16*** | F : 5´- **aagctt**TAAGTTGATAAAGCTAATTTC -3´<br>R : 5´- **ggtacc**TTTGGGAGGGAATTAAAGGG -3´ |
| ***pNsCBTS2α*** | F : 5´- **aagctt**AACTATGAAAAAATTTTAAC -3´<br>R : 5´- **ggtacc**CTCTCTCTTTCTCTCGCCAAAC -3´ |
| ***pNtLTP1*** | F : 5´- **aagctt**GGGCAGACAAAATTAG -3´<br>R : 5´- **ggtacc**TCTTAAAAGAAAAATTA -3´ |
| ***pNtCPS2*** | F : 5´- **aagctt**CTGCAAATCTCCCAACATTATC -3´<br>R : 5´- **ggtacc**TTTCTAATTTAATTTTTGTTTTATTC -3´ |
| ***pNtRbcS-T1*** | F : 5´- **gcggccgc**TCTGGTCTCGACCTTGATG -3´<br>R : 5´- **ggtacc**GTTCACCTTGACTTTCAACTAC -3´ |
| ***pNtMALD1*** | F : 5´- **gcggccgc**ACTAAATTTTGATTACTTTAAAACTGTGG -3´<br>R : 5´- **ggtacc**TTTTTTTTTCCTCTAAAAAAACTTGGAGTG -3´ |
