## Supplemental Figures for "Identification of two new trichome-specific promoters of *Nicotiana tabacum*"

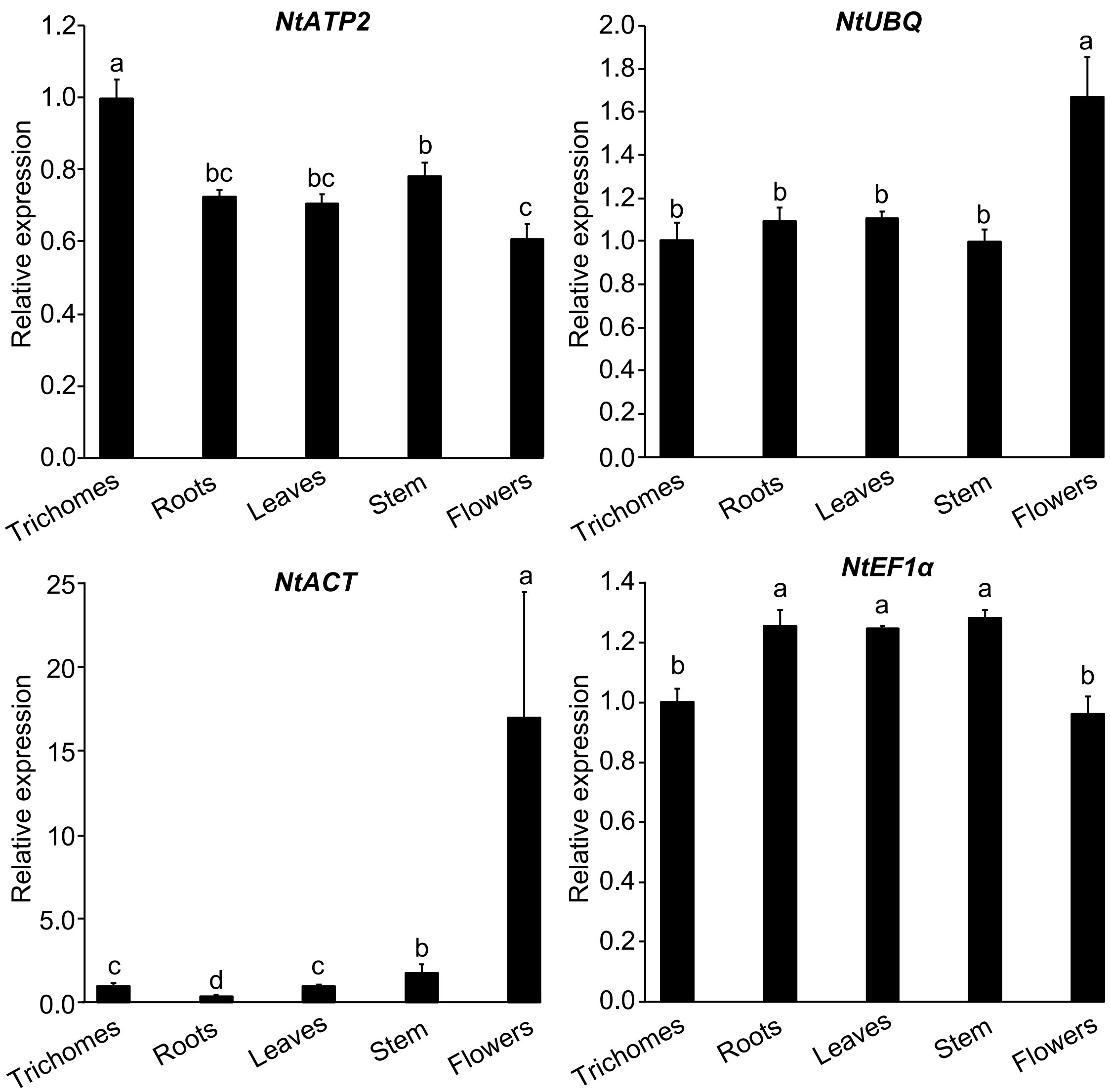


**Supplemental Fig. 1** Transcript levels of control genes in different organs of *N. tabacum*. *NtATP2*, *NtUBQ,* and Nt*EF1α* control genes were used to normalize the data in Figure 1. *NtACT* gene was not selected for normalization due to its instability. Transcript levels were normalized to the geometric mean of those of *NtATP2, NtUBQ,* and *NtEF1α* genes. Results are shown as mean ± SD of three to five repeats. Different letters indicate significant differences according to a Kruskal-Wallis test ($p < 0.05$) followed by a Tukey post hoc test

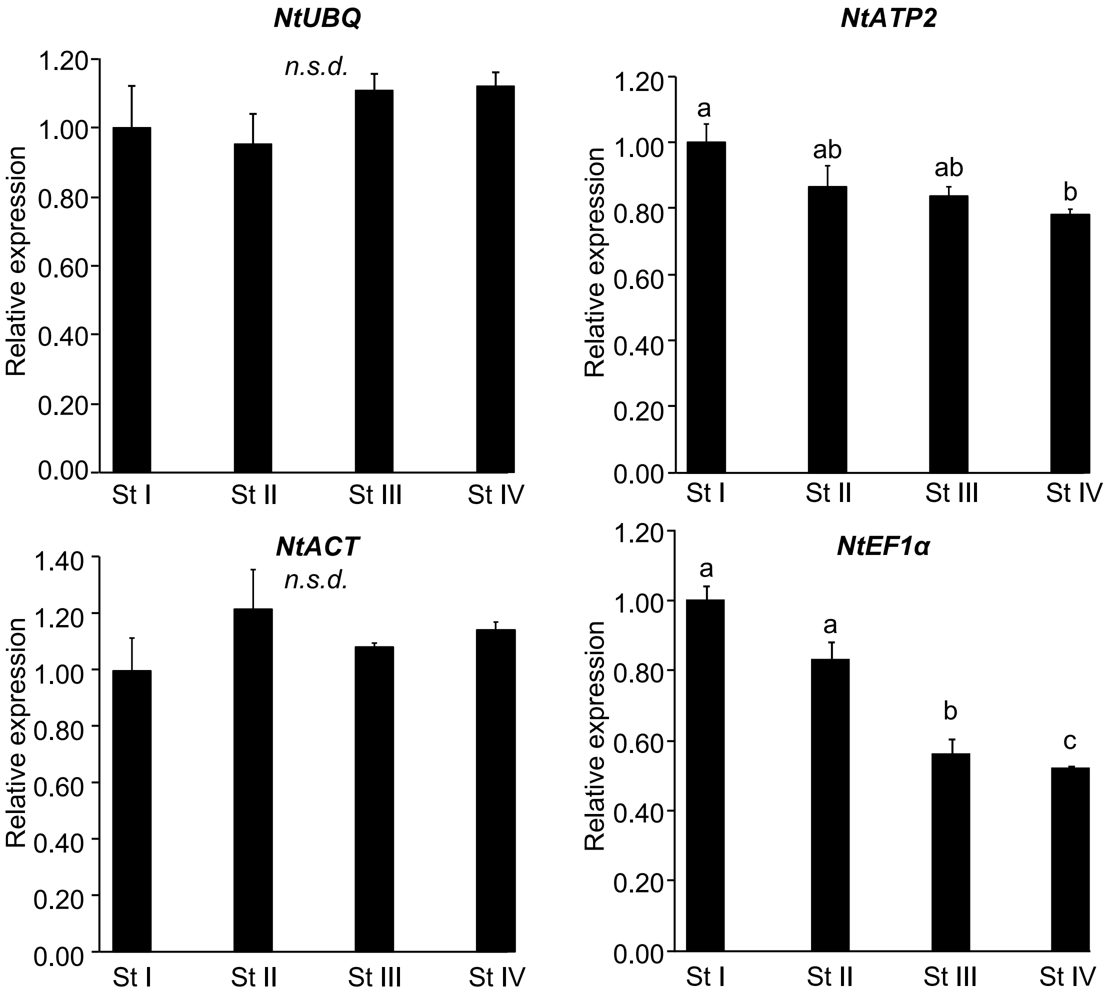


**Supplemental Fig. 2** Transcript levels of control genes in trichomes isolated from N. tabacum leaves at different developmental stages. *NtUBQ, NtATP2, and NtACTIN* control genes were used to normalize the data in Figure 2. *NtEF1α* gene was not selected for normalization due to its large decrease in transcript levels in trichomes during leaf development. Transcript levels were normalized to the geometric mean of those of *NtUBQ*, *NtATP2*, and *NtACTIN* genes. St: leaf developmental stage. Stage 1: leaf length < 2.5 cm; stage II: leaf length between 2.5 cm and 6.5 cm; stage III: leaf length between 6.5 cm and 15 cm; stage IV: leaf length > 15 cm. Results are shown as mean ± SD of three repeats. Different letters indicate significant differences according to a Kruskal-Wallis test ($p < 0.05$) followed by a Tukey post hoc test. *n.s.d.*: no significant differences

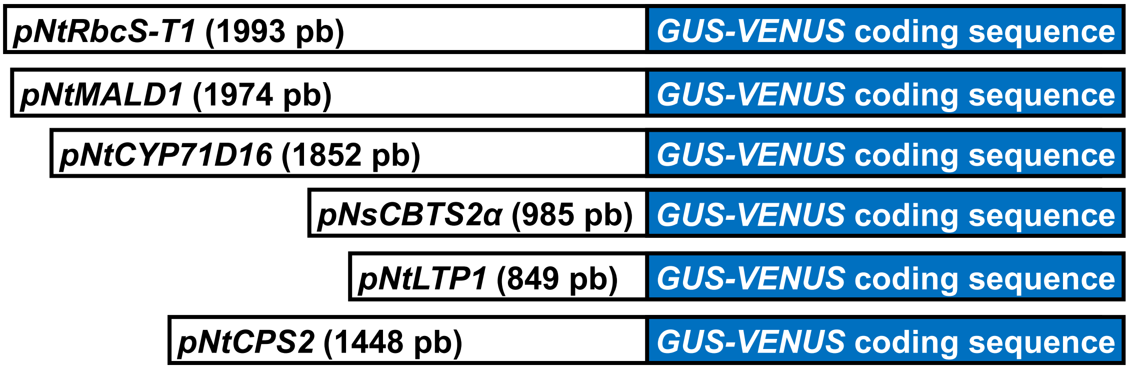


**Supplemental Fig. 3** Molecular constructs used to generate transgenic *N. tabacum* expressing the *GUS-VENUS* reporter gene under the control of trichome-specific promoters. The transcription promoter regions of *NtRbcS-T1* (MG493459.1), *NtMALD1* (MG493458.1), *NsCBTS2α* (HM241151.1), *NtLTP1* (AB625593.1), *NtCYP71D16* (AF166332.1), and *NtCPS2* (HE588139.1) were amplified and cloned as described in the Materials and methods

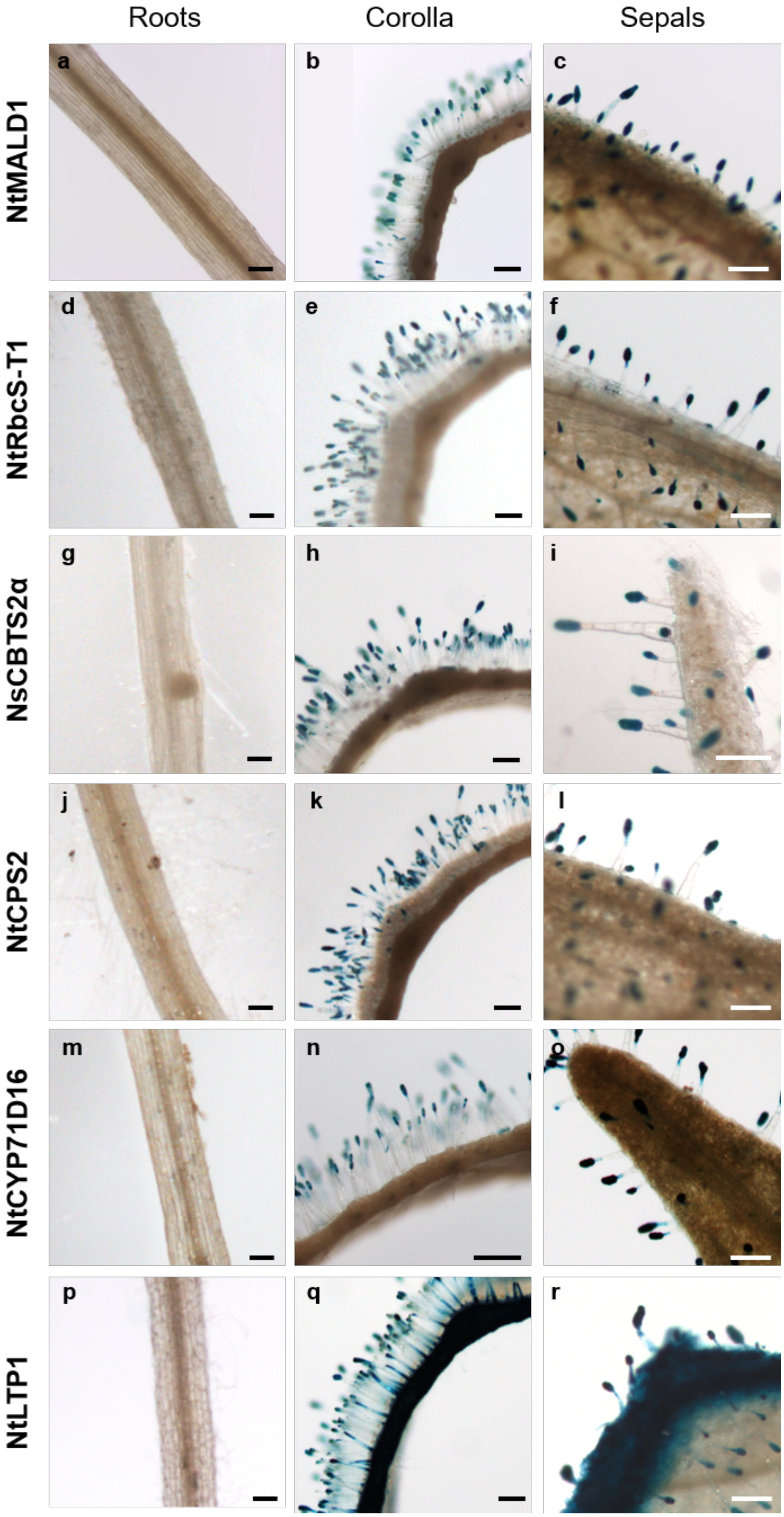


**Supplemental Fig. 4** GUS activity in various tissues of *N. tabacum* transgenic lines. GUS staining was performed on transgenic lines expressing *GUS-VENUS* under the control of the promoter region of *NtMALD1* (**a**-**c**), *NtRbcS-T1* (**d**-**f**), *NsCBTS2α* (**g**-**i**), *NtCPS2* (**j**-**l**), *NtCYP71D16* (**m**-**o**), and *NtLTP1* (**p**-**r**). While no GUS signal was detected in roots, a trichome specific signal was detected in corolla and sepals for all constructs but the *NtLTP1 r*eporter that displayed a wider expression pattern. Scale bars: 200 µm
